# Intracellular FMRpolyG-HSP70 complex: Possible use as biochemical marker of Fragile X Tremor Ataxia Syndrome

**DOI:** 10.1101/278416

**Authors:** Giuseppe Bonapace, Rosa Gullace, Daniela Concolino, Grazia Iannello, Radha Procopio, Monica Gagliardi, Gennarina Arabia, Gaetano Barbagallo, Angela Lupo, Lucia Ilaria Manfredini, Grazia Annesi, Aldo Quattrone

## INTRODUCTION

Fragile X-associated tremor/ataxia syndrome (FXTAS) is a late-onset neurodegenerative disorder that affects about 25% of carriers of non-coding CGG-repeat expansions (55–200 repeats) in the premutation range within the fragile X gene (*FMR1*). Main clinical features include intention tremor, cerebellar ataxia, and Parkinsonism [1]. Recently, great emphasis on the pathogenetic role of toxic aggregates, produced by a RAN translation process on the 5’ expanded CGG region, has been given [2,3]. These aggregates contain a small protein with a polyglycin stretch, (FMRpolyG), and so far, have been isolated and characterized in drosophila, mouse model and post mortem FXTAS patients biopsies, but never in living patients. [4] The early diagnosis of this syndrome, in adult carriers of premutations, remains difficult due to the absence of specific markers and, the complexity of the procedure aimed at producing human cellular models, makes the study of the role of these aggregates in the pathogenesis of FXTAS difficult as well. [5]

Here we demonstrate, for the first time, using immunocytochemistry (ICC) and Western Blot procedure (WB), that it is possible to easily assess the presence of aggregates containing FMRpolyG protein, in fibroblasts isolated from a clinical and molecular characterized FXTAS living patient, and discuss some possible practical implications.

## MATERIALS AND METHODS

### Case report

The patient was a 50-year male with a history of tremor and slowness since age 43. He also complained of occasional syncopal episodes. His past and family histories were unremarkable. Neurological examination revealed broad-based gait, facial hypomimia and mild asymmetric (right greater than left) resting tremor in the distal upper and lower extremities, in association with postural and action tremor in the upper limbs. Tone, strength, and sensation were normal. There were also a slight bradykinesia and increased deep tendon reflexes. General physical examination was normal, without sexual or urinary disturbances, except for orthostatic hypotension with a decrease in systolic and diastolic blood pressure of 34 and 14 mmHg respectively, changing position from supine to standing upright. A generalized cerebral atrophy with severe involvement of the cerebellum and of the middle cerebellar peduncles were present in 3T MRI.

### Gene Scan analysis

Amplicons depending on the size of the CGG repeat region, were produced using a specific FRAXA Triple Primed PCR Kit (Abbott Molecular). Hi-Di formamide, (Applied Biosystems), and 2μl of ROX 350 size standard (Celera Corporation) were combined with 2μl of PCR product. Samples were denatured at 95°C for 2 minutes before loading onto an ABI 310 with POP-6 polymer on a 36-cm or 50-cm array. Data were analyzed using GeneMapper v.3.7 software.

### Immunocytochemistry

Patient’s and non FXTAS premutated control (60 repeats NFPC) fibroblasts from skin biopsies, were plated in chamber slide (Nunc Rochester NY) and incubated 48 h in RPMI 10% FBS. After paraformaldehyde fixation, permeabilization with 0.2% Triton X-100 followed by blocking in 0,2% gelatin at 4°C, cells were incubated for 3h at room temperature with a anti FMRpolyG monoclonal Antibody (clone 9FM 1-B7 Millipore USA) and a goat anti Human HsP70 1:200 (Santa Cruz). Anti-mouse FITC conjugated and anti-goat TRITC conjugated specifc secondary antibody 1:500 were used for the detection. Hundred microscopic fields were analyzed and cells containing aggresomes (positive) were counted and evaluated as percentage of the total cells.

### Immunoprecipitation studies

After precleaning with 60 μl of a Staphylococcus A membrane, the fibroblast’ cytosolic fraction, was isolated and incubated overnight at 4°C with a Rabbit anti human HsP70 antibody. The immunocomplexes were separated by SDS page and Western blot and analyzed by using a Mouse anti human FMRpolyG and a goat antimouse HRP conjugated. In a separate experiment, the same rabbit anti Hsp70 antibody, with a secondary anti rabbit HRP conjugated, was used to assess the amount of free Hsp70 (without immunoprecipitation) in fibroblasts from NFPC and patient.

### Statistical Analysis

A Kruskall-Wallis test was used to evaluate significant differences in the number of cell positive for FMRpolyG-HSP70 complex, between NFPC and patient.

## RESULTS

The molecular analysis of the FMR1CGG repeats shows in our patient a mosaicism for a 44 and 116 CGG expansions in the premuation range for FRAXA. (Fig 1A).

**Fig 1A.**
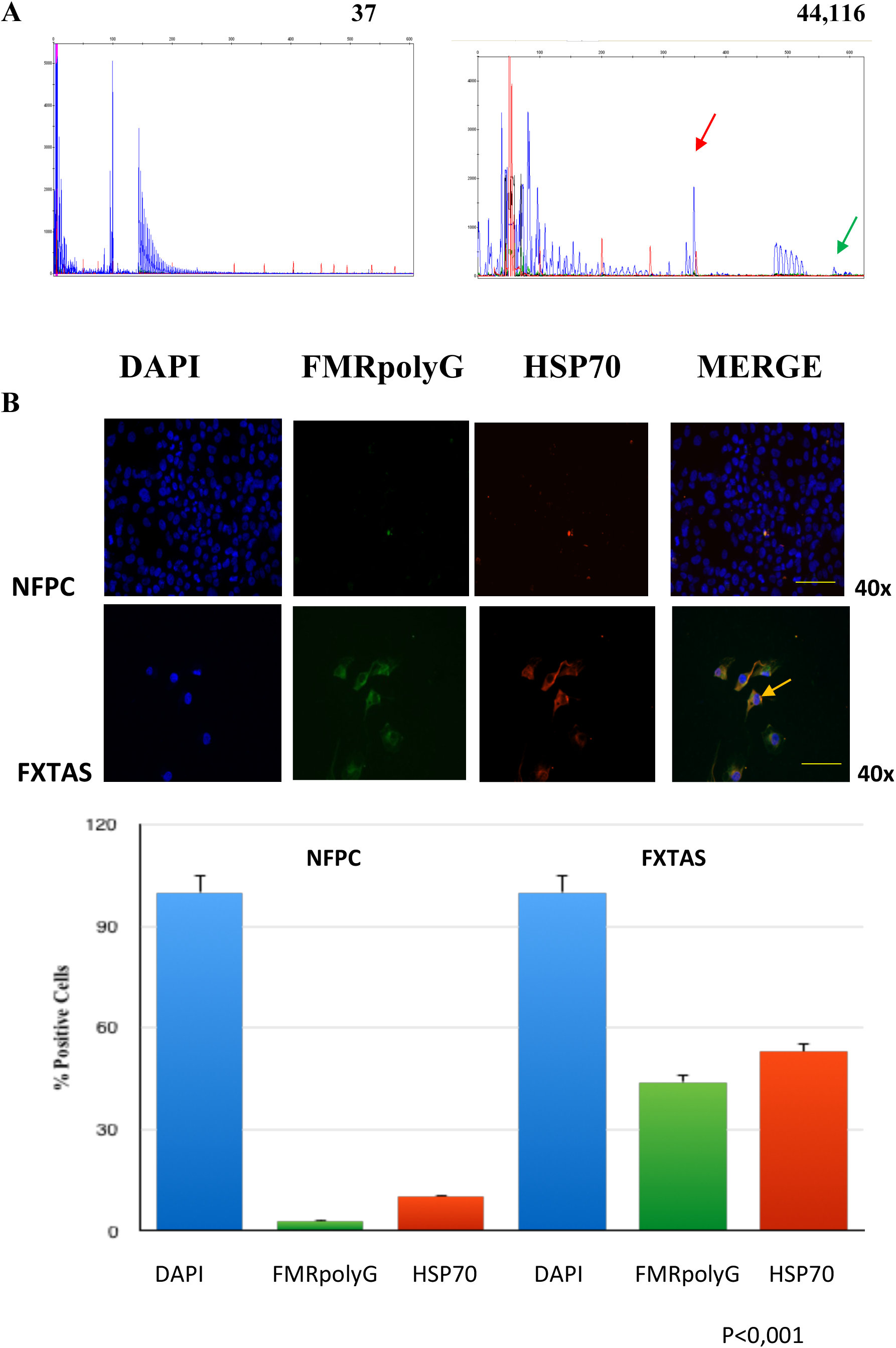
Genescan analysis of the CGG repeats within FMR1 gene from Normal male control (Left) and from FXTAS patient (Right): Alleles size are shown in the upper right corner. The patient was a mosaic for a 44 allele 1(red arrow) and for a 116 allele (green arrow) B **Recruitment of HSP70 cellular chaperone to FMRpolyG:** Fibroblasts from FXTAS patient and NFPC were processed for indirect immunofluorescence using antibodies against FMRpolyG and against HSP70. HSP70 and FMRpolyG colocalize in about 45% of FXTAS fibroblasts No association is evident in NFPC (Non FXTAS premutated control)

To assess the production of FMRpolyG protein we performed immunocytochemistry experiments both on FXTAS patient’s and NFPC fibroblasts.

Fig 1B shows that the FMRpolyG accumulates in patient’s fibroblasts both in the cytoplasm and nucleus (yellow arrow). As expetcted in presence of soluble aggregates, there is a clear colocalization of FMRpolyG with the Heat Shock protein 70 (HSP70), a well known cellular chaperone involved in the protection against the effects triggered by aggregosomes. (See Supplemental Information.)

To demonstrate that the HSP70 does not merely colocalize with FMRpolyG, but it is a component of the same intracellular complex involved in a defensive reaction against the aggregates, we performed immunoprecipitation experiment on the cytosolic fibroblasts fraction.

Fig 2A shows that the anti HSP70 antibody is able to immunoprecipitate the FMRpolyG protein, confirming that there is a specific physical interaction with FMRpolyG as described in the first step of proteosomal mediated degradation pathway. [6]

**Fig 2.**
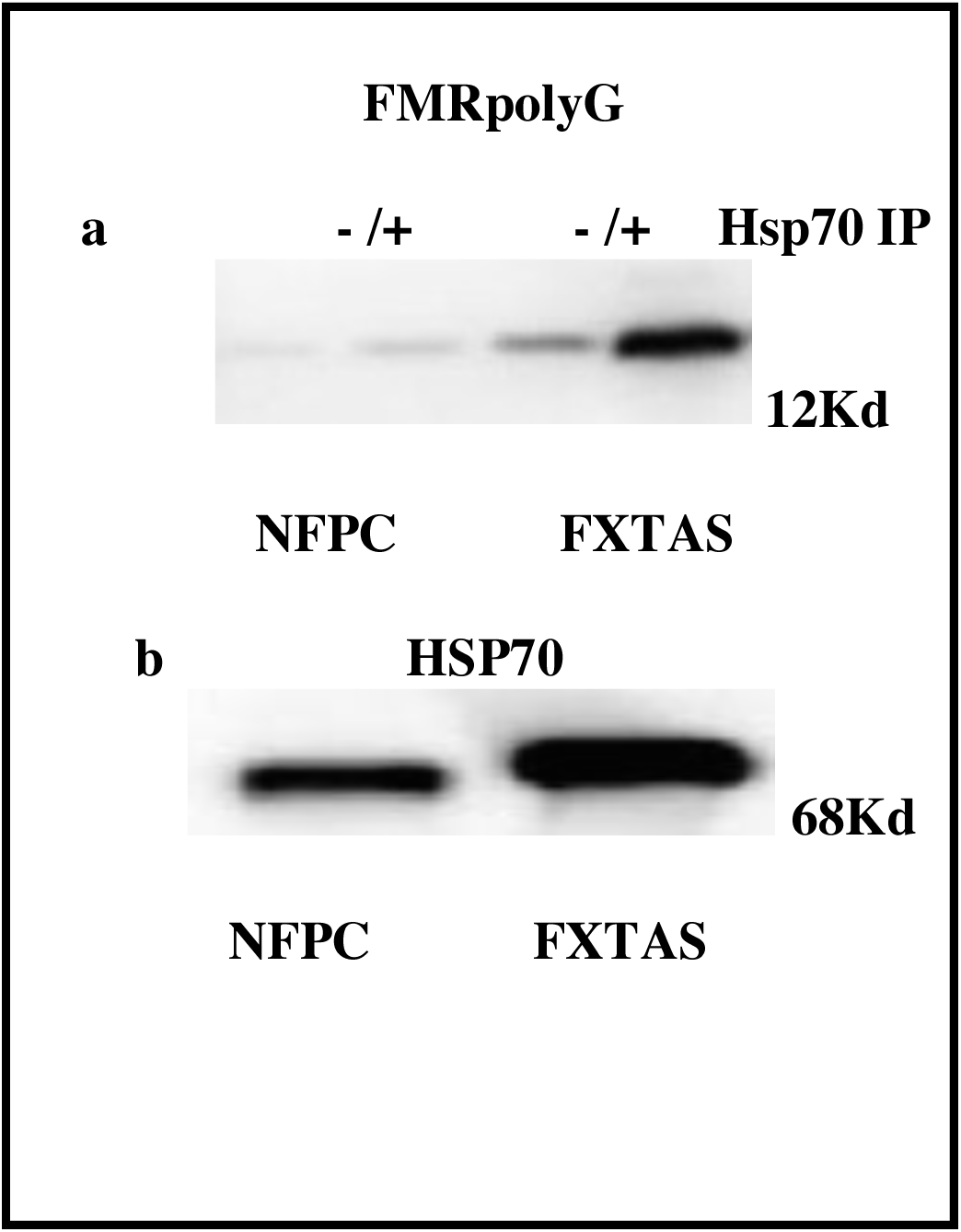
Western Blot Analysis of HSP70 FMRpolyG interaction: **(a)** Cellular extracts from patient’s fibroblast and NFPC, immunoprecipitated with anti HSP70 were separated on 12% polyacrilammide gel. After transfer, the filter was incubated with an anti FMRpolyG Ab and developped with a secondary HRP conjugated antibody. **(b)** Western blot of HSP70 protein on the same amount of patient and NFPC cellular extracts. Strikingly, the recruitment of HSP70 by FMRpolyG and the increased levels of total HSP70 in the patient’s fibroblasts are confirmed +IP: Immunoprecipitation -IP: No immunoprecipitation

Western blot only for the HSP70 protein on the same amount of cellular extract is shown in figure 2B. According to the defense response, the HSP70 levels are increased in the FXTAS fibroblasts in comparison to the steady state levels in NFPC.

## DISCUSSION

Features of FXTAS overlap many other neurodegenerative disorders, including Parkinson, Alzheimer diseases and frontotemporal dementia (e.g., progressive cognitive impairment, altered mood and behavior). [7] These observations underscore the need to develop a new and reliable diagnostic tool, to confirm the diagnosis of FXTAS. Todd and colleagues recently demonstrated, on postmortem biopsies and murine models, as a translation of CGG repeats without ATG (RAN) occurs in two out of the three frames, giving rise to short proteins containing either a polyalanine or a polyglycine stretch and that polyglycine protein resulted in the formation of heterogeneous protein inclusions, which were toxic both in neuronal transfected cells and in Drosophila. [3]

Based on this observation we asked if would have been possible to assess the presence of such aggregates in vivo, in order to open the way towards a novel diagnostic marker for FXTAS in living patients. Starting from the observation that the FMR1 protein is expressed in a wide range of cells, [8] we demonstrate, the presence of FMRpolyG aggregates in fibroblasts from a living patient carrier of a premutated FMR1 allele associated to clinical signs of FXTAS but not in a normal premutated carrier of CGG expansion. To our knowledge this is the first observation in living patients.

Because the increase in the HsP70 level has been described as marker of cellular defense against the inclusions found in neurons and astrocytes from FXTAS post mortem brain, [9], we also studied such reaction in the fibroblasts of our patient. Our data demonstrating an increase in the amount of total Hsp70, along with an evident molecular interaction between the FMRpolG peptide and the HsP70, confirm also in our patient, the presence of a protective cellular response against such peptide. Even though the validation of this procedure needs larger number of FXTAS patients to be evaluated in order to assess its diagnostic and predictive value, these data demonstrate that it is possible to assess the presence of potential toxic FMRpolyG-HSP70 aggregates in skin biopsies from FXTAS living patients, establish a new simple, fast and reproducible human cellular model to study in *vivo* the progression of the disease and suggest that in a close future this procedure can be exploited to confirm the clinical diagnosis in patients showing neurological signs of FXTAS associated to a FMR1 CGG premutated allele.

## AKNOWLEDGMENTS

### Author Contributions

**Giuseppe Bonapace**, Study concept and design, immunohistochemistry and WB analysis

**Rosa Gullace**, Cell Colture and WB experiments

**Daniela Concolino**, Analysis and interpretation of data

**Grazia Iannello**, Molecular Biology data

**Radha Procopio**, Molecular Biology data

**Monica Gagliardi**, Fragment analysis

**Grazia Annesi**, Study concept and design, critical revision of manuscript

**Gennarina Arabia**, Draft paper, clinical data collection

**Gaetano Barbagallo**, Draft paper and skin biopsy

**Angela Lupo**, Clinical data collection

**Lucia Ilaria Manfredini**, Clinical data collection

**Aldo Quattrone**, Study supervision, Critical revision of manuscript for intellectual content

